# AICAR-mediated selection of karyotypically normal cells in Pallister-Killian syndrome patient-derived skin fibroblasts

**DOI:** 10.1101/2022.01.27.478025

**Authors:** Emanuela Salzano, Maninder Kaur, Anastasia M Jacko, Deborah McEldrew, Sarah E Raible, Ian D Krantz, Kosuke Izumi

## Abstract

Pallister-Killian Syndrome (PKS) is a mosaic aneuploidy syndrome typically caused by the presence of a supernumerary marker isochromosome containing two extra copies of the short arm of chromosome 12 (iso-chromosome 12p or tetrasomy 12p). This isochromosome is always present in a mosaicism state that has tissue limited distribution. PKS is characterized by severe neurodevelopmental delay, intellectual disability, multisystem involvement and congenital malformations including typical dysmorphic features and skin pigmentation anomalies.

Aneuploid cells, irrespective of the identity of the supernumerary chromosome, including cancer cells, yeast cells and mouse embryonic fibroblasts (MEFs), have been demonstrated to present a disruption of protein homeostasis and increased basal stress levels; resulting in a greater sensitivity to chemical compounds inducing cellular energy stress compared to euploid cell lines. The burden of trisomy 21 has also been recently shown to impair the proteostasis network in lymphoblastoid cell lines and fibroblasts obtained from individuals with Down syndrome.

In this study, we demonstrate that AICAR, 5-Aminoimidazole-4-carboxamide ribonucleotide, a known energy stress inducing drug with antiproliferative effects on aneuploidy cancer cells and MEFs, is also able to selectively eliminate cells carrying the isochromosme12p in PKS clones in a time and dosage dependent manner. Collectively, our results indirectly provide evidence of increased basal energy and proteotoxic stress in PKS cells carrying isochromosome 12p, and suggest a potential therapeutic drug-based strategy that, selectively acting as a stressor for aneuploid cells, may establish the euploid state in PKS and a broader spectrum of human mosaic disorders.

## Introduction

Pallister-Killian Syndrome (PKS, MIM 601803) is a mosaic aneuploidy syndrome typically caused by the presence of a mosaic supernumerary marker isochromosome 12p. PKS is characterized by typical craniofacial findings, skin pigmentary anomalies, neurodevelopmental delay, intellectual disability and several congenital malformations [Izumi et al., 2014]. Other than medical management, pharmacologic-based therapeutic approaches for PKS have not previously been proposed.

Due to the mosaic nature of PKS, only a certain percentage of cells in an individual with PKS have the isochromosome. It is believed that the euploid cells in individuals with PKS arose from cells that have lost the isochromosome (since the origin of the isochromosome is in meiosis, the original zygote would have had this marker). The euploid cells are suggested to have a growth advantage over the aneuploid cells and further skew the mosaic ratio towards the euploid cell lines [Conlin et al., 2012]. Therefore, approaches that are able to selectively eliminate aneuploid cells could represent a therapeutic strategy.

Aneuploidy syndromes generally cause severe problems in human development and growth and have previously been considered irremediable. Recently, several *in vitro* methods, collectively referred as “Chromosome Therapies”, have been proposed as a potential treatment for aneuploid syndromes. They result in loss, silencing or replacement of aberrant chromosomes in cells *in vitro*. Strategies for a new potential therapy may include molecule and drug treatments able to induce selection or promote a euploid state in cells [Plona et al. 2016].

Since aneuploidy is often seen in cancer cells, the mechanistic insights for correcting chromosomal abnormalities can be extrapolated from cancer research studies. In studies evaluating the efficacy of several energy-stress and/or proteotoxic compounds, many have demonstrated AICAR (5-Aminoimidazole-4-carboxamide ribonucleotide) to be one of the most effective as a potential anticancer agent for aneuploidy driven cancers [Neckers and Neckers, 2002; Manchado and Malubres et al. 2011; Tang et al.2011; Ly et al., 2013]. AICAR treatment has also been found to selectively inhibit proliferation of MEFs trisomic for chromosomes 1, 13, 16 and 19, in a dosage dependent manner [Tang et al. 2011]. AICAR is a cell permeable precursor of ZMP (an AMP analog) which allosterically activates AMP-activated protein kinase (AMPK), mimicking cellular energy stress [Corton et al. 1995].

The purpose of this study is to evaluate the efficacy of AICAR in diminishing the number of cells carrying the isochromosome 12p in mosaic samples of PKS and to indirectly assess effects on their underlying stress replicative mechanism and its ability to covert constitutional mosaic clones to the euploid state.

## Materials and Methods

### Ethical compliance

All PKS probands were enrolled under an institutional review board approved protocol at The Children’s Hospital of Philadelphia.

### Cell Culture

Immortalized and primary skin fibroblasts cell lines from probands with PKS and normal controls (or wild type cell line - WT) at early passages (<p15) were uniformly cultured in RPMI 1640 (Life Technologies, Carlsbad, CA, USA) supplemented with 15% FBS (Hyclone South Logan, UT, USA), antibiotics (100 units/ml of penicillin and 100 ug/ml of streptomycin), and 1% L-glutamine (Life Technologies, Carlsbad, CA, USA). Cells were incubated at 37 °C in a humidified atmosphere of 5% C02. Cells were trypsinized (0.25% trypsin-EDTA) and split at 75%-80%confluency. We established hTERT-immortalized cell lines from fibroblasts derived from probands with PKS as described previously [Izumi et al. 2015].

### Drug Treatments

Approximately 1.5×10^3^ euploid and PKS immortalized/primary fibroblasts have been respectively plated into 24-well-plates and, after 24 hours were exposed to AICAR [Toronto Research Chemicals, Toronto, ON, Canada]. Different doses of AICAR (from 0.2mM, up to 1.5mM concentration) were tested for a variable time up to 13 days of treatment. Experiments were repeated in triplicate.

### Cell viability assay, DNA isolation and quantification of mosaic level by digital droplet PCR (ddPCR)

The effect of the drug has been evaluated both through cell viability using the trypan blue viability assay (Sigma-Aldrich) and through mosaic ratio quantification by ddPCR Technology (Bio-rad’s QX100tm/Copy Number Assay) at different indicated time points. The mosaic ratio of iso12p cells was quantified as described previously [Fujiki et al. 2016]. DNA samples were extracted from PKS skin fibroblasts and normal control skin fibroblasts using the Puregene DNA isolation kit (Qiagen, Valencia, CA) with modified protocol. DNA yields and quality were determined using the Nanodrop 2000 Spectrophotometer (#E112352; Thermo Scientific, Somerset, NJ).

A total of 30-60 ng of genomic DNA was used to perform ddPCR analysis. FAM fluorescent tagged TaqMan^®^ copy number assay probes detecting the *ETNK1* gene (Hs01860263_cn) (Life Technologies, Carlsbad, CA) was used to quantify the copy number of 12p. The ddPCR system was operated according to the manufacturer’s instruction. The PCR reaction solution was compartmentalized by a droplet generator, and then, PCR amplification performed. After the PCR amplification was completed, a droplet reader counted the number of droplets that were positive and negative for FAM and VIC fluorophore. The mosaic ratio was calculated based on the total positive signal counts of *ETNK1* normalized against the TERT positive signal counts.

### Statistical Analysis

All data are shown as the mean of three independent experiments ± standard deviation. Data analysis were performed and plotted using the Prism software package, version 6. A 2-way Anova with Sidak’s multiple comparisons test has been used to test differences in mosaic ratio and among relative cell number means between euploid and PKS cell lines in absence and after different dosages of drug treatment. A p-value of < 0.05 was deemed to be significant (*p-value < 0.05; ** p-value < 0.01; *** p-value < 0.001; **** p-value <0.0001).

## Results

### Cellular proliferation profile of PKS fibroblasts

Since recent studies have demonstrated that gain of extra chromosomes in yeast [Torres et al., 2007], mouse embryonic fibroblasts (MEFs) [Williams et al., 2008] and human Down syndrome (DS) fibroblasts [Kimura et al., 2005] leads to slowed proliferation *in vitro* as the result of an associated stress response to the aneuploid status, we initially examined the proliferation profile of cultured PKS cells of approximately 50% mosaic ratio. Both WT and PKS clones were plated at the same initial density and counted by the trypan blue assay at common time points. The proliferation ability of primary PKS fibroblasts was reduced as compared to the immortalized PKS cells (Fig. 1), likely due at least in part to the higher mitotic index itself of immortalized cell lines. However, both immortalized and primary PKS clones showed a significantly impaired proliferation compared to those of control cells. (Fig. 1A/B).

**Fig. 1.**
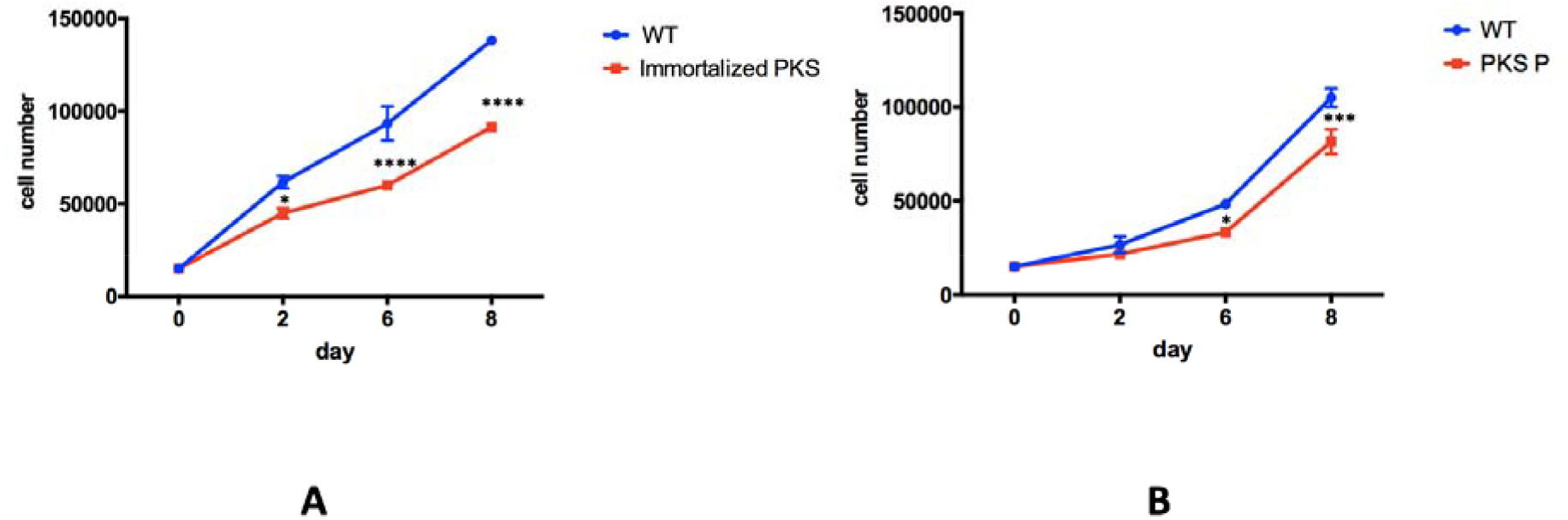
Cellular Proliferation Profile of PKS fibroblasts. Growth curves of immortalized (A) and non-immortalized primary (B) PKS fibroblasts and wild-type control cells. The cell lines were tested in triplicates (* p value <0.05, *** p-value < 0.001****p value <0.0001; 2-way ANOVA multiple comparison).

### AICAR effects on immortalized PKS fibroblasts

To verify if PKS immortalized clones were preferentially sensitive to AICAR, cells were cultured with different drug concentrations and evaluated at different time points assessing for any morphological changes, reduction in cell number and level of mosaicism. Due to the mosaic nature of the PKS clones, we hypothesized that we would detect a mosaic ratio reduction if the drug acted selectively on aneuploid cells. To test for AICAR effects, we started treating the cells with high AICAR dosages of 1mM and 1.5mM. Then, we progressively decreased the concentration of the drug to identify a dose that would be effective in PKS fibroblasts and safe for normal controls.

Overall, cell number counts showed that immortalized euploid fibroblasts (WT) tolerate the compound better than PKS fibroblasts. Morphological changes and cell number reduction of treated immortalized PKS cell line were detectable under optic microscopy after 48 hours of drug exposure and proportionally correlated with AICAR doses and time of exposure (fig.2).

**Fig. 2.**
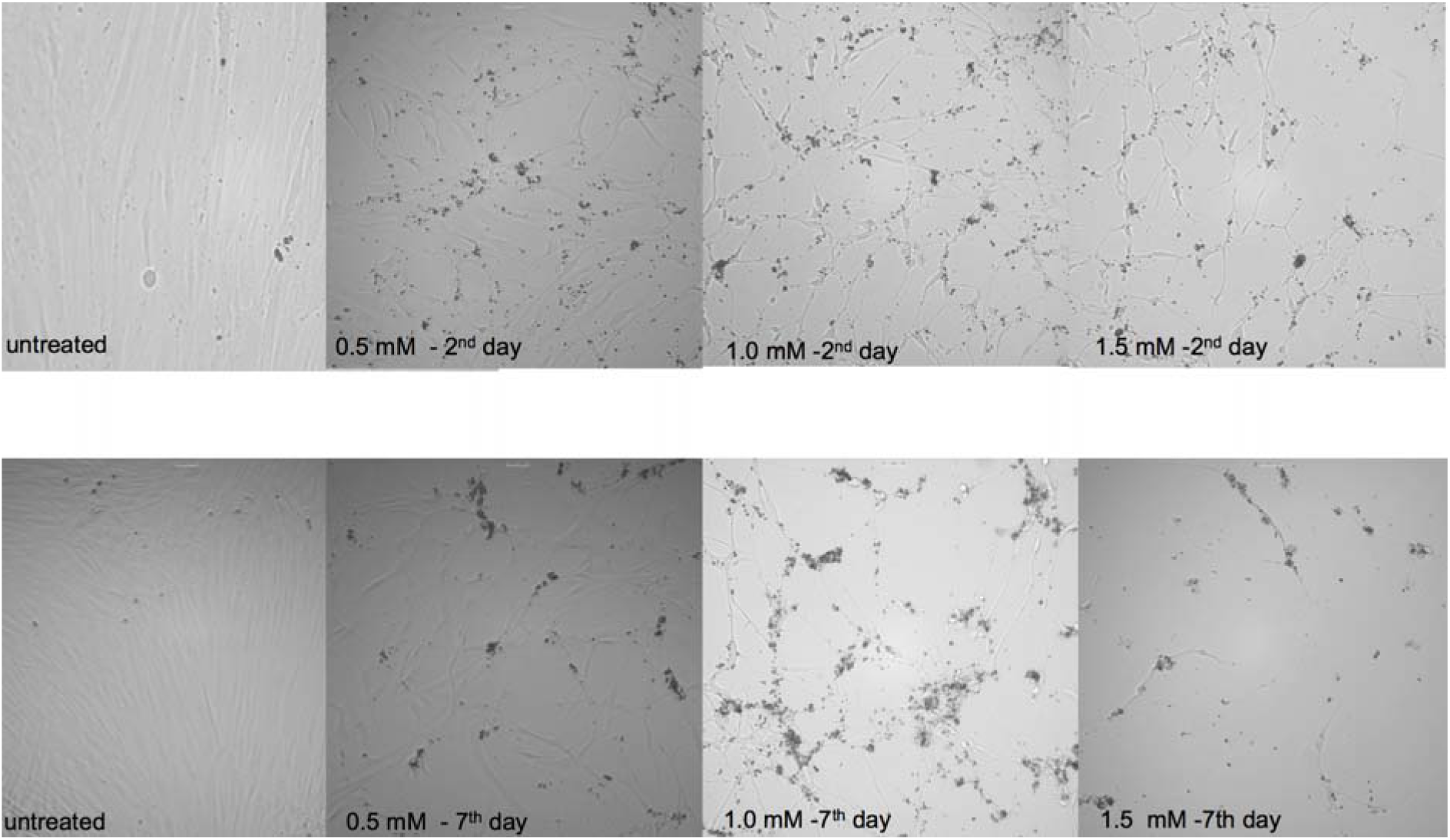
AICAR inhibits Proliferation in immortalized PKS fibroblasts by the 2^nd^ day of treatment. Morphological changes in immortalized PKS fibroblasts under optic microscope respectively after 2 and 7 days of 0.5 mM, 1mM 1.5 mM AICAR exposure.

The effects of AICAR treatment on PKS fibroblasts number was confirmed to be proportional to the dosage (R=0.672) (fig. 3).

**Fig. 3.**
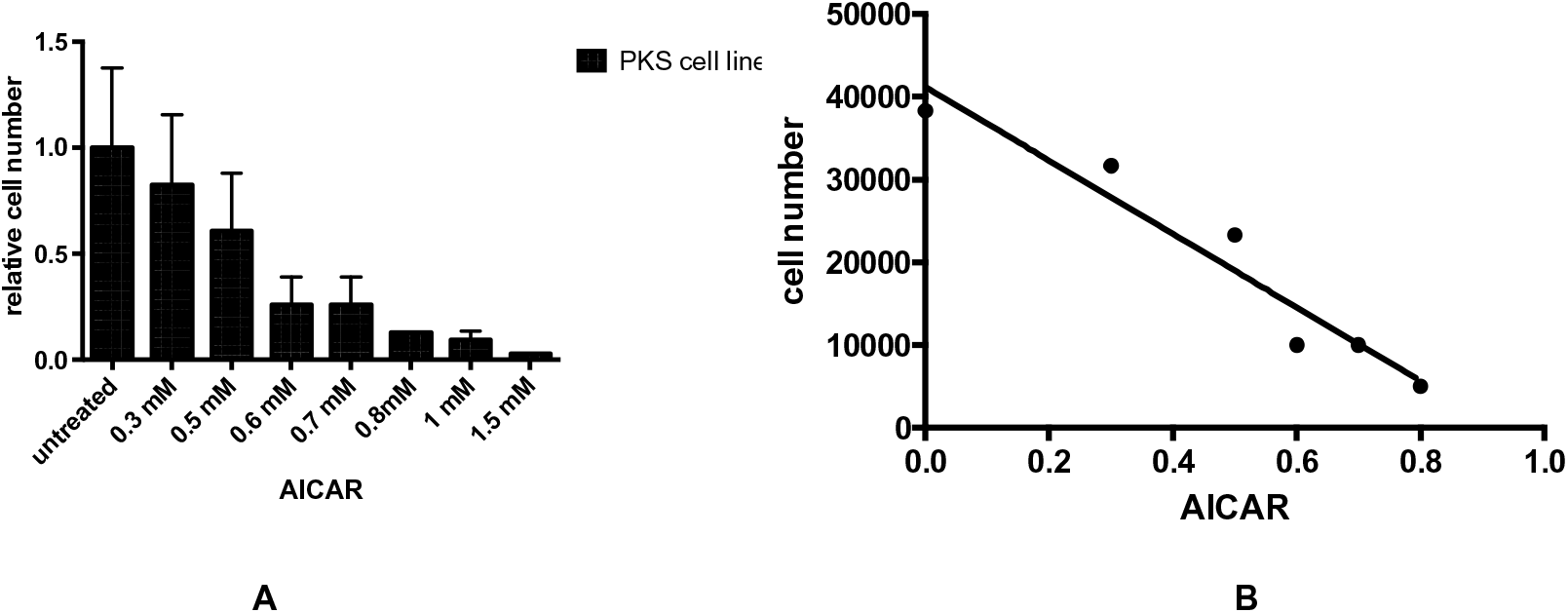
AICAR dosage – dependent effects: Immortalized PKS fibroblasts were exposed to different AICAR concentrations **A)** Cell number as relative percentage of the untreated control was determined after 7 days of drug treatment. **B)** Linear Regression Analysis Chart (R square 0.6720; p value < 0.0001). Data are the mean of three independent experiments ± standard deviation.

To specifically assess the cut-off safety dose for control cells and verify if any density dependent effect was detectable, experiments were performed by plating both wild type euploid cells and PKS clones at different initial densities and counting cells after 2 and 13 days of treatment (fig.4).

**Fig. 4.**
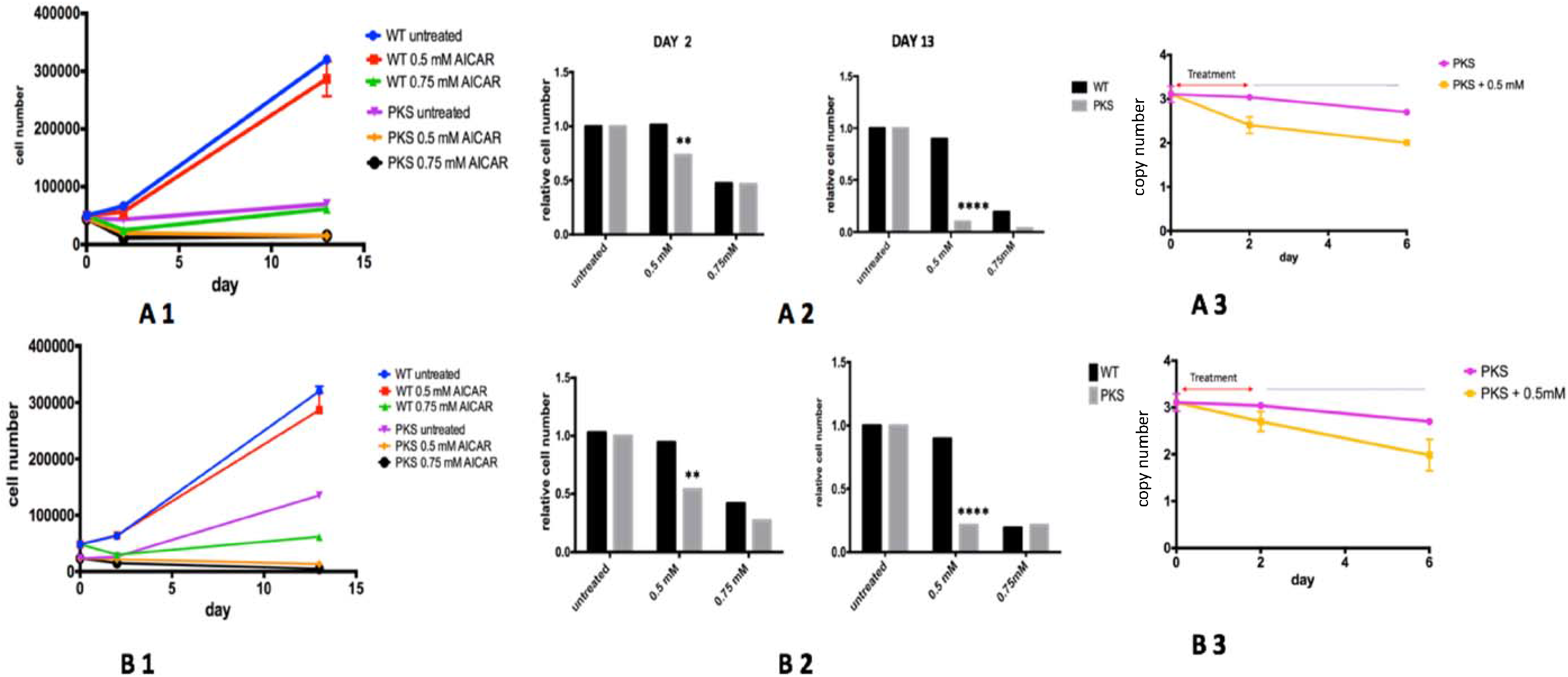
AICAR selectively inhibits proliferation and reduces mosaic ratio of immortalized PKS clones. Wild Type euploid (WT) and PKS fibroblasts (PKS) were plated at different initial density respectively of 1.25×10^^4^ × 1.9 cm^2^ **(A)** and 2.5×10^^4^ **(B)**. They both were grown in absence (blue WT/purple PKS) or presence of two different AICAR concentrations (red WT/yellow PKS 0.5 mM treatment; green WT/black PKS 0.75 mM treatment). 1. Cell growth curves: cell number was determined after 2 days of treatment and after 13 days of treatment. 2. Cell number as relative percentage of the untreated control after 2 days and 13 days of treatment. The 0.5 mM AICAR dosage was able to selectively inhibit the PKS clone proliferation (** p value <0.005, ****p value <0.0001; 2-way ANOVA multiple comparison) by acting in a density cellular density dependent manner. **3**. The mosaic level variation has been reported as mosaic ratio changes assessed by ddPCR evaluated at 48 hours of exposure, and after 4 days of drug discontinuation. 48 hours of AICAR exposure at 0.5 mM dosage induced a significant mosaic ratio reduction in PKS fibroblasts. Reduction of mosaic level was inversely proportional to cell density (A3 p value <0.0001, B3 p value<0.005). Y-axis represents 12p copy number.

Although an AICAR concentration ≥ 0.75 mM was toxic for both PKS and euploid fibroblasts, a dosage of 0.5 mM was able to significantly reduce the number of PKS immortalized fibroblasts after 2 days of treatment (p <0.005, 2-way ANOVA multiple comparisons test) without affecting the control cell line, even when long term exposure, up to 13 days, was performed (p < 0.0001, 2-way ANOVA multiple comparisons test) (fig. 4 A.1/2-B.1/2).

After 13 days of drug exposure, the number of fibroblasts from PKS probands was significantly lower than those from controls (p<0.0001, 2-Way ANOVA multiple comparisons test), irrespective of the initial cell density. Also, at the same dosage and time of exposure, cellular death rate resulted in a 46% increase among cells plated at initial 1.25 × 10^3^ density compared to that of fibroblast clones initially plated at double the cell density of 2.5 ×10^3^ (p value < 0.0001, 2-Way ANOVA multiple comparisons test). This suggests a cellular density-dependent drug effect (p value < 0.0001, 2-Way ANOVA multiple comparisons test) (fig.4 A2/B2).

The mosaic ratio in PKS skin fibroblasts was found to be stable and close to a euploid state after 5 days of continued AICAR treatment at 0.5 mM dosage. However, 2 days of 0.5 mM AICAR exposure was sufficient to reach a significant mosaic ratio reduction that spontaneously tended to euploidy within the following 4-5 days after drug discontinuation and remained steady at least for 2-3 more days. (fig.4 A3/B3). An initial lower cell density with the same drug dosage of 0.5mM allowed for a more significant mosaic ratio reduction after 2 days of treatment (p<0.0001, 2-Way ANOVA test) compared to experiments performed with double cell density (p<0.005, 2 Way ANOVA test) (fig.4 A3/B3).

### AICAR effects on primary PKS fibroblasts

Having characterized the effects of AICAR treatment on immortalized PKS skin fibroblasts, we wanted to determine whether the same effects were reproducible on primary skin fibroblasts from PKS patients. AICAR effects on primary cell lines were evaluated using the same methodology applied for immortalized skin fibroblasts. Primary PKS fibroblasts were generally more sensitive to AICAR exposure compared to normal control cell lines and morphological changes were detectable under optic microscopy after 48 hours of drug exposure. Similarly, to what we observed in immortalized PKS cell lines, the quantitative reduction of cells in the treated primary PKS cell line was confirmed to proportionally correlate with AICAR dosage (R= 0.8671; p value <0.0001) (fig. 5A). The drug concentration of 0.5 mM was able to differentially act on cellular growth of PKS clones without affecting the normal cell line after 2 days and to induce a significant reduction in primary PKS fibroblasts after 6 days of treatment (fig. 5B). The mosaic ratio reduction in primary PKS clones was observed after 8 days of 0.5 mM AICAR exposure (p <0.05), reaching a value closer to the euploid state after 6 days of drug discontinuation (p <0.0001) (fig. 5C). Although the same trend in reduction of tetrasomic cells in primary PKS clones was obtained in half the time by using a concentration of 1mM (p <0.05), high AICAR dosage > 0.7 mM tended to result in more toxicity (for both the PKS and WT cell lines), and was especially pronounced for treatments lasting longer than 72 hours.

**Fig. 5.**
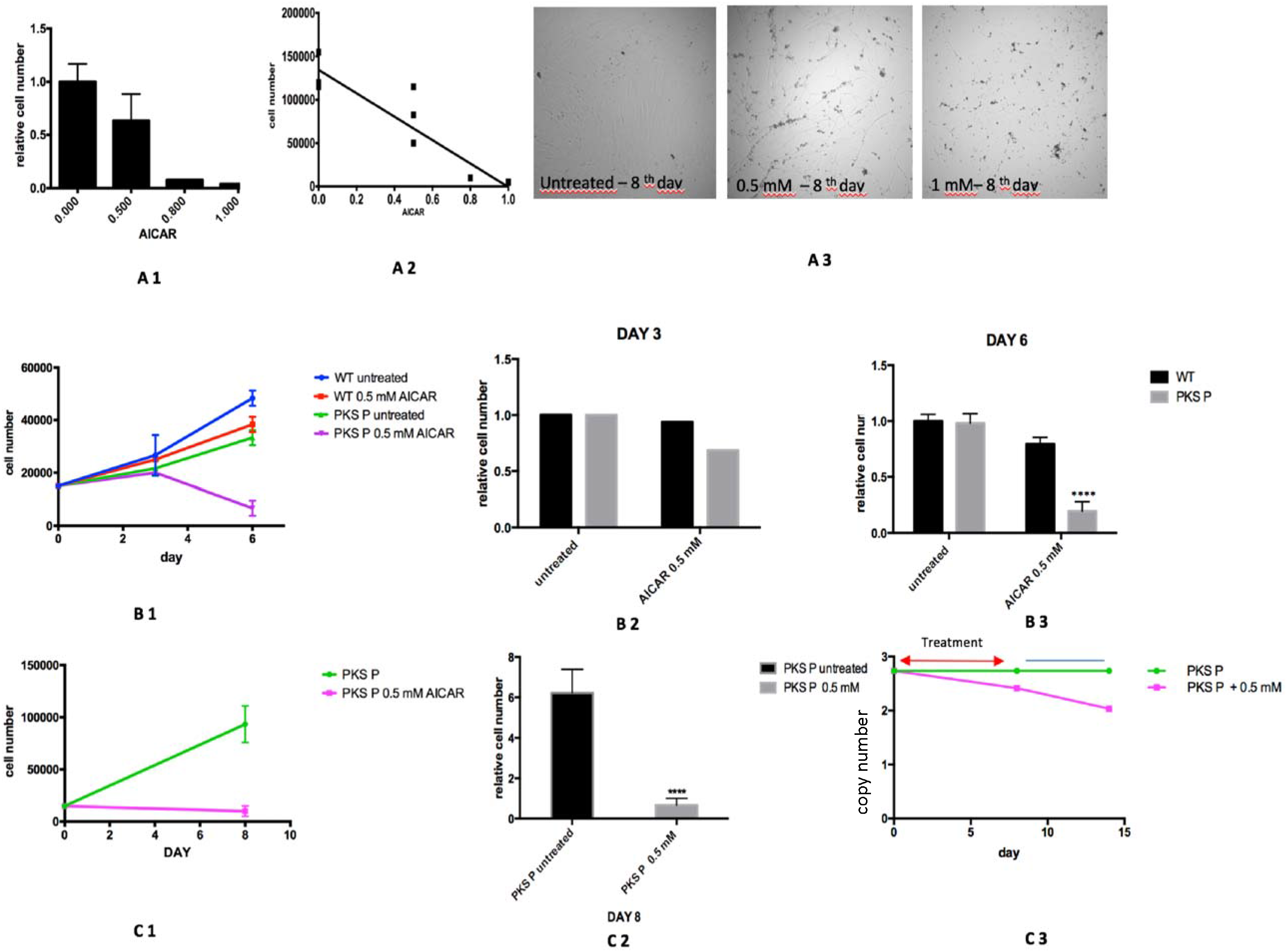
AICAR inhibits proliferation of primary PKS clones. **A) AICAR dosage-dependent effect**. Primary PKS fibroblasts were exposed to different AICAR concentrations **A1)** Cell number as relative percentage of the untreated control was determined after 8 days of drug treatment as the percentage of the untreated control **A2)** Linear Regression Analysis Chart (R 0.8671; p value <0.0001) **A3)** Morphological changes under optic microscope respectively after 8 days of 0.5 mM and 1mM AICAR exposure. **B) 0.5 mM AICAR effects on cell number**. Wild Type euploid cells (WT) and non-immortalized primary PKS fibroblasts (PKS P) were grown in absence (blue WT/green PKS P) or presence of 0.5 mM AICAR concentration (red WT/purple PKS P). **B1)** Cell growth curves. Cell number was determined after 3 days of treatment **(B2)** and after 6 days of treatment **(B3)** (****p value <0.0001; 2-way ANOVA multiple comparison). AICAR induces a significant cell number reduction after 6 days of treatment **C) 0.5 Mm AICAR effect after 8 days of treatment**. Cell Growth curves of treated PKS clones compared to untreated PKS clones **(C1)**. Significant PKS cell number reduction was demonstrated after 8 days of drug exposure (**C2) (**\*\*\*\*p value < 0.0001, 2-way ANOVA). The mosaic level variation has been reported as mosaic ratio changes assessed by ddPCR: it has been reported after 8 days of exposure, and after 6 days of drug discontinuation. AICAR was able to selectively inhibit proliferation of aneuploid fibroblasts in primary PKS clones after 8 days of treatment (p<0.05, 2 Way ANOVA test) getting close to the euploid status after 6 days of drug discontinuation (p <0.0001, 2 Way ANOVA test) **(C3)**. Data are the mean of three independent experiments ± standard deviation. Y-axis represents 12p copy number.

## Discussion

Using *in vitro* cellular models, we have demonstrated that AICAR can reduce the percentage of PKS aneuploid cells. Our data clearly indicates that both immortalized and primary PKS skin fibroblast cell lines are more sensitive to AICAR than controls. Although primary PKS cell lines required longer exposure, intermediate concentrations of AICAR are able to selectively eliminate aneuploidy in both immortalized and primary cell lines carrying the isochromosome 12p.

The molecular mechanism by which AICAR reduced the PKS aneuploid cells remains unknown. However, based on the previous studies, we hypothesize that a dysfunctional proteostasis, secondary to the presence of the isochromosome 12p, may be involved in an increased basal cellular stress in the tetrasomic fibroblasts, making them more sensitive to the drug.

Systematic studies have shown that the presence of an additional chromosome elicits a set of shared metabolic phenotype changes in both trisomic yeast cells and mouse embryonic fibroblasts (MEFs) and in aneuploid cancer cells [Torres et al., 2007; De Berardinis et al., 2007; Williams et al., 2008; Tang et al. 2011; Oromendia et al. 2012]. Irrespective of the identity of the supernumerary chromosome, aneuploid cells activate protein degradation and folding pathways in an attempt to correct protein stoichiometry imbalances caused by the aneuploidy [Torres et al., 2007; De Berardinis et al., 2007; Williams et al., 2008]. When folding load exceeds cellular protein quality-control machinery capacity, proteotoxicity induces upregulation of stress-response genes and down-regulation of cell proliferation genes (environmental stress response (ESR)-like signature), increasing factors (aneuploidy-associated protein signature (APS)) involved in the response to oxidative stress and leading to impaired proliferation or to apoptosis by p53 activation [Donnelly et al., 2014; Oromendia and Amon 2014; Santaguida and Amon 2015]. Conversely, this results in an increased sensitivity of aneuploid cells to increased proteotoxic and energy stress compounds compared to normal controls [Torres et al., 2007; De Berardinis et al., 2007; Williams et al., 2008; Santaguida and Amon 2015]. Different studies have also suggested that aneuploidy is associated with premature aging and reduced lifespan [Baker et al. 2004; Baker et al., 2008, Wijshake et al., 2012; Baker et al., 2013].

As far as we know, Down Syndrome (DS) has been the only *in vitro* human model investigated among constitutional aneuploidies, confirming a disrupted proteostasis network (including unfolded protein response, chaperone system and proteasomal degradation), and an enhanced loss of viability compared to euploid cell lines when treated with ER stressors [Westerheide et al., 2005; Anckar et al., 2011; Aivazidis et al. 2017].

The *in vitro* cellular proliferation growth profile of PKS clones was impaired compared to those of karyotypically normal clones, strongly suggesting that the presence of the small extra chromosome may be responsible for an underlying dysfunctional energy and proteotoxic stress-response. Analysis of previously published expression profiles of primary fibroblasts from PKS patients [Kaur et al. 2014], shows that there are several differentially expressed genes known to be involved in proteostasis networks and cell cycle regulation (including *CCND2, COP7A, USP5A, UFDL1, DEPDC6, HSPA2, HSPA12A*). Consistent with the identification of a defective heat shock protein response and reduced action of the HSP70 pathway in the *in vitro* DS cellular model [Aivazidis et al. 2017], both the *HSPA2* and *HSPA12A* genes are significantly downregulated in PKS clones. *HSPA2*, in particular, plays a pivotal role in the protein quality control system, ensuring the correct folding of proteins, the re-folding of misfolded proteins and targeting of proteins for subsequent degradation. Also, the *DEPDC6* gene (more commonly known as *DEPTOR*) which functions as a negative modulator of both the mTorc1 and mTorc2 signaling pathways, has been found to be downregulated in PKS cells compared to controls cell line. It is interesting to note that recent studies have also highlighted that a bidirectional crosstalk exists between ER stress and mTor signaling; as cell toxicity by ER stress is coupled to the perpetual activation of mTORC1 that in turn promotes apoptosis by the Ire1a–ASK1–JNK pathway [Appellnzer-Herzog et al. 2012; Catena et al.2017; Kato et al. 2012].

AICAR is an adenosine analog taken up into cells by adenosine transporters and phosphorylated by adenosine kinase, thus generating the AMP-mimetic, AICAR monophosphate (ZMP) [Corton et al., 1995; Kim et al., 2016]. ZMP can accumulate in millimolar concentrations in cells, directly activate AMPK and, being a natural intermediate in the purine nucleotide synthetic pathway, be metabolized by AICAR transformylase, which catalyzes synthesis of the purine nucleotide inosinate. However, as an AMP analog, AICAR activates many other AMP-dependent enzymes and acts as a “cross-road” for multiple pathways, including mToR signaling, that modulates cellular growth, apoptosis, autophagy, cellular aging, lipogenesis, ROS generation and DNA repair [Ly et al. 2013; Liu et al. 2014]. Based on these data, AICAR likely triggers apoptosis in tetrasomic 12p fibroblasts due to an exaggerated stress-response [Tang et al. 2011].

Because of the favorable physiological outcomes of AMPK activation on metabolism, AMPK has been considered to be an important therapeutic target for controlling human diseases, including metabolic syndromes. AMPK has been shown to be activated by any modulator that causes AMP or calcium accumulation, including numerous naturally occurring compounds and phytochemicals (eg. polyphenols, alfa-lipoic acid, ginsenoside, salicylate) [Kim et al. 2016]. Thus, further investigations into the molecular mechanisms mediating AICAR’s effects on aneuploidy cells would be useful to identify additional compounds acting on the same pathways but potentially without the adverse effects of a chemotherapeutic compound.

There are some potential limitations to this study. One challenge is the mosaic nature of PKS itself, which complicates all cell based *in vitro* studies and may limit the possibility to translate the use of the drug into clinical practice. However, as far as we know, this is the first human constitutional mosaic model treated by an anti-proliferative drug to rescue cellular clones from aneuploidy.

Although more studies are needed to assess AICAR’s molecular effects on different structural mosaicisms, our results provide a cellular level characterization of AICAR’s effect and of the underlying replicative stress mechanism in a PKS cellular model. Our data may also offer a new way of understanding tetrasomy 12p and it would be important to determine whether chronic proteotoxic cellular stress may contribute deleteriously to the phenotype. While a critical region on 12p13.3 has been defined that likely contains important genes in driving certain aspects of the PKS phenotype, [Izumi et al.., 2012], the proteotoxic burden secondary to mosaic aneuploidy levels in brain tissue itself might induce a neurodegenerative process and contribute to a common neuro-developmental phenotype among human structural aneuploidies [Oromendia et al., 2014].

In conclusion, we have demonstrated that intermediate/high dosages of AICAR are able to selectively eliminate fibroblasts carrying the isochromosome 12p and help maintain a euploid state in PKS clones. As the effect of the drug appears to be dosage and cellular density dependent, it is also possible to define a potentially safe drug dosage level, raising the interesting possibility for new treatment strategies for PKS as well as a broader spectrum of human mosaic chromosomal disorders.

## Acknowledgements

We are deeply indebted to the individuals with PKS and their families, who participated in this study, as well as to the PKS family support groups for their continuous contribution. The project has been made possible by Grants of PKS Kids Italia Onlus Fondation and PKS Kids USA Foundation (ES, KI) and CHOP CdLS and Related Diagnoses Center Endowment Funds (IDK, SR). We would also like to thank the Italian Society of Pediatrics who supported ES with the SIP “Pioneers of Pediatrics” Award.

## Notes

### Competing Interest Statement

The authors have declared no competing interest.

### Summary of Updates

"Pallister Killian syndrome" was corrected to "Pallister-Killian syndrome" in the title.

## References

Aivazidis S, Coughlan CM, Rauniyar AK, Jiang H, Liggett LA, Maclean KN, Roede JR. The burden of trisomy 21 disrupts the proteostasis network in Down syndrome. PLoS One. 2017 Apr 21;12(4): e0176307.

Anckar J, Sistonen L. Regulation of HSF1 function in the heat stress response: implications in aging and disease. Annu Rev Biochem. 2011; 80:1089–115.

Appenzeller-Herzog C, Hall MN. Bidirectional crosstalk between endoplasmic reticulum stress and mTOR signaling. Trends Cell Biol. 2012;22(5):274–82.

Baker DJ, Jeganathan KB, Cameron JD, Thompson M, Juneja S, Kopecka A, Kumar R, Jenkins RB, de Groen PC, Roche P, van Deursen JM. BubR1 insufficiency causes early onset of aging-associated phenotypes and infertility in mice. Nat Genet. 2004;36(7):744–9.

Baker DJ, Perez-Terzic C, Jin F, Pitel KS, Niederländer NJ, Jeganathan K, Yamada S, Reyes S, Rowe L, Hiddinga HJ, Eberhardt NL, Terzic A, van Deursen JM. Opposing roles for p16Ink4a and p19Arf in senescence and ageing caused by BubR1 insufficiency. Nat Cell Biol. 2008;10(7):825–36.

Baker DJ, Dawlaty MM, Wijshake T, Jeganathan KB, Malureanu L, van Ree JH, Crespo-Diaz R, Reyes S, Seaburg L, Shapiro V, Behfar A, Terzic A, van de Sluis B, van Deursen JM. Increased expression of BubR1 protects against aneuploidy and cancer and extends healthy lifespan. Nat Cell Biol. 2013;15(1):96–102.

Catena V, Fanciulli M. Deptor: not only a mTOR inhibitor. J Exp Clin Cancer Res. 2017 Jan 13;36(1):12.

Conlin LK, Kaur M, Izumi K, Campbell L, Wilkens A, Clark D, Deardorff MA, Zackai EH, Pallister P, Hakonarson H, Spinner NB, Krantz ID. Utility of SNP arrays in detecting, quantifying, and determining meiotic origin of tetrasomy 12p in blood from individuals with Pallister–Killian syndrome. Am J Med Genet A 2012;158A:3046–3053.

Corton JM, Gillespie JG, Hawley SA, Hardie DG. 5-aminoimidazole-4-carboxamide ribonucleoside. A specific method for activating AMP-activated protein kinase in intact cells? Eur J Biochem. 1995; 229: 558–565.

DeBerardinis RJ, Mancuso A, Daikhin E, Nissim I, Yudkoff M, Wehrli S, Thompson CB. Beyond aerobic glycolysis: transformed cells can engage in glutamine metabolism that exceeds the requirement for protein and nucleotide synthesis. Proc Natl Acad Sci U S A. 2007;104(49):19345–50.

Donnelly N, Passerini V, Dürrbaum M, Stingele S, Storchová Z. HSF1 deficiency and impaired HSP90-dependent protein folding are hallmarks of aneuploid human cells. EMBO J. 2014;33(20):2374–87.

Fujiki K, Shirahige K, Kaur M, Deardorff MA, Conlin LK, Krantz ID, Izumi K. Mosaic ratio quantification of isochromosome 12p in Pallister-Killian syndrome using droplet digital PCR. Mol Genet Genomic Med. 2016;4(3):257–61.

Izumi K, Nakato R, Zhang Z, Edmondson AC, Noon S, Dulik MC, Rajagopalan R, Venditti CP, Gripp K, Samanich J, Zackai EH, Deardorff MA, Clark D, Allen JL, Dorsett D, Misulovin Z, Komata M, Bando M, Kaur M, Katou Y, Shirahige K, Krantz ID. Germline gain-of-function mutations in AFF4 cause a developmental syndrome functionally linking the super elongation complex and cohesin. Nat Genet. 2015; 47, 338–344.

Izumi K, Krantz ID Pallister–Killian syndrome. Am J Med Genet C Semin Med Genet. 2014;166C:406–13.

Kato H, Nakajima S, Saito Y, Takahashi S, Katoh R, Kitamura M. mTORC1 serves ER stress-triggered apoptosis via selective activation of the IRE1–JNK pathway. Cell Death Differ. 2012; 19, 310–320.

Kaur M, Izumi K, Wilkens AB, Chatfield KC, Spinner NB, Conlin LK, Zhang Z, Krantz ID. Genome-wide expression analysis in fibroblast cell lines from probands with Pallister Killian syndrome. PloS one. 2014; 9(10), e108853.

Kim J, Yang G, Kim Y, Kim J, Ha J. AMPK activators: mechanisms of action and physiological activities. Exp Mol Med. 2016;48(4):e224.

Kimura M, Cao X, Skurnick J, Cody M, Soteropoulos P, Aviv A. Proliferation dynamics in cultured skin fibroblasts from Down syndrome subjects. Free Radic Biol Med. 2005;39(3):374–80.

Liu X, Chhipa RR, Pooya S, Wortman M, Yachyshin S, Chow LM, Kumar A, Zhou X, Sun Y, Quinn B, McPherson C, Warnick RE, Kendler A, Giri S, Poels J, Norga K, Viollet B, Grabowski GA, Dasgupta B. Discrete mechanisms of mTOR and cell cycle regulation by AMPK agonists independent of AMPK. Proc Natl Acad Sci U S A. 2014;111(4):E435–44.

Ly P, Kim SB, Kaisani AA, Marian G, Wright WE, Shay JW. Aneuploid human colonic epithelial cells are sensitive to AICAR-induced growth inhibition through EGFR degradation. Oncogene. 2013;32(26):3139–46.

García Martínez J, García-Inclán C, Suárez C, Llorente JL, Hermsen MA. DNA aneuploidy-specific therapy for head and neck squamous cell carcinoma. Head Neck. 2015;37(6):884–8.

Manchado E, Malumbres M. Targeting aneuploidy for cancer therapy. Cell. 2011;144(4):465–6

Plona K, Kim T, Halloran K, Wynshaw-Boris A. Chromosome therapy: Potential strategies for the correction of severe chromosome aberrations. Am J Med Genet C Semin Med Genet. 2016;172(4):422–430.

Neckers L, Neckers K. Heat-shock protein 90 inhibitors as novel cancer chemotherapeutic agents. Expert Opin Emerg Drugs. 2002; 7, 277–288

Oromendia AB, Dodgson SE, Amon A. Aneuploidy causes proteotoxic stress in yeast. Genes Dev. 2012;26(24):2696–708.

Oromendia AB, Amon A. Aneuploidy: implications for protein homeostasis and disease. Dis Model Mech. 2014;7(1):15–20.

Santaguida S, Amon A. Short- and long-term effects of chromosome mis-segregation and aneuploidy. Nat Rev Mol Cell Biol. 2015;16(9):576.

Tang YC, Williams BR, Siegel JJ, Amon A. Identification of aneuploidy-selective antiproliferation compounds. Cell. 2011;144(4):499–512.

Torres EM, Sokolsky T, Tucker CM, Chan LY, Boselli M, Dunham MJ, Amon A. Effects of aneuploidy on cellular physiology and cell division in haploid yeast. Science. 2007;317(5840):916–24.

Westerheide SD, Morimoto RI. Heat shock response modulators as therapeutic tools for diseases of protein conformation. J Biol Chem. 2005;280(39):33097–100.

Wijshake T, Malureanu LA, Baker DJ, Jeganathan KB, van de Sluis B, van Deursen JM. Reduced life-and healthspan in mice carrying a mono-allelic BubR1 MVA mutation. PLoS Genet. 2012;8(12):e1003138.

Williams BR, Prabhu VR, Hunter KE, Glazier CM, Whittaker CA, Housman DE, Amon A. Aneuploidy affects proliferation and spontaneous immortalization in mammalian cells. Science. 2008;322(5902):703–9.

